# Late I_Na_ activation of cardiac TTX-sensitive sodium channels by AaH-II induces an arrhythmogenic phenotype

**DOI:** 10.1101/2025.09.17.676709

**Authors:** Hugo Millet, Pauline Belhumeur, Agnès Hivonnait, Floriane Bibault, Matthias Dereli, Agnès Tessier, Morteza Erfanian, Barbara Ribeiro, Flavien Charpentier, Mikael Croyal, Jérôme Montnach, Michel De Waard

## Abstract

**Aims:** Late sodium current (I_NaL_) is a key contributor to cardiac arrhythmias, but its precise origin and arrhythmogenic potential from tetrodotoxin-sensitive (TTX-S) sodium (Na_v_) channels remain unclear. While the FDA-endorsed toxin ATX-II has been widely used to model I_NaL_-associated arrhythmogenesis, it lacks selectivity, limiting its utility in dissecting the roles of individual Na_v_ channel subtypes. This study investigates the proarrhythmic impact of TTX-S Na_v_ channel activation using AaH-II*, a scorpion venom-derived peptide with selective efficacy for TTX-S channels.

**Methods and Results:** Using automated patch-clamp recordings, we characterized AaH-II* selectivity across human Na_v_ isoforms and demonstrated potent, preferential activation of I_NaL_ in hNa_v_1.1, 1.2, 1.3, and 1.6 over the TTX-resistant cardiac isoform hNa_v_1.5. Calcium imaging in isolated adult rat cardiomyocytes showed that low nanomolar concentrations of AaH-II* induced spontaneous calcium release events and arrhythmogenic calcium transients, even in the absence of Na_v_1.5 activation. *Ex vivo* multielectrode array recordings in Langendorff-perfused rat hearts confirmed dose-dependent ventricular conduction slowing, prolonged repolarization, and increased arrhythmia burden, all mitigated by TTX. *In vivo*, intravenous AaH-II* administration in rats elicited QTc prolongation, atrioventricular block, and ventricular tachyarrhythmias, which were significantly suppressed by TTX pretreatment.

**Conclusion:** We identify AaH-II* as a powerful and selective tool to study I_NaL_ from TTX-S Na_v_ channels in cardiac tissue. Our findings reveal that TTX-S channel-mediated I_NaL_ alone is sufficient to induce arrhythmias and that pharmacological inhibition of these channels offers a promising antiarrhythmic strategy. These results advocate for broader consideration of TTX-S Na_v_ channels as targets in arrhythmia research and drug safety screening.

## Introduction

In the heart, action potential (AP) upstroke is supported by the opening of voltage-gated Na^+^ channels, which in turns triggers the opening of voltage-gated Ca^2+^ channels required to ignite muscle contraction through ryanodine receptor recruitment and sarcoplasmic reticulum (SR) Ca^2+^ release. The fine-tuning of this entire process is required for the perfect synchronization of cardiomyocyte contraction in the cardiac tissue. This adjustment is made possible thanks to the precise time-dependent activation of voltage-gated K^+^ channels that terminate the AP. Not surprisingly, an imbalance in the normal kinetic behavior of any one of these cardiac ion channels will lead to arrythmia. In the case of LQT3, *SCN5A* variants produce Na_v_1.5 sodium currents with abnormally prolonged inactivation kinetics manifested by a plateau phase (usually called late I_Na_ (I_NaL_)) ^1^. The consequences of this altered channel kinetic are a prolonged AP plateau that produce a delayed pro-arrhythmic repolarization. Besides rare genetic causes, arrythmia may also develop in patients with heart failure, a condition that favors the appearance of I_NaL_. Finally, I_NaL_ may also inadvertently be sparked by drugs which explains why the Comprehensive in vitro Proarrhythmia Assay (CiPA) procedure includes a stringent control on this issue. In spite of its pathophysiological importance, I_NaL_ is particularly difficult to investigate without artificially augmenting it since it represents only a tiny fraction of the total cardiac Na^+^ current (< 0.1%). To better apprehend the role of I_NaL_, the use of the sea anemone toxin II (ATX-II), a natural promoter of this late Na^+^ current has been advocated by the Federal Drug Administration (FDA). The pro-arhythmic consequences of promoting I_NaL_ by ATX-II have been deciphered and include prolonged activation of Ca^2+^ channels, enhanced intracellular Na^+^ concentration, SR Ca^2+^ overload, CaMKII activation, leaky ryanodine receptors, enhanced NCX Na^+^/Ca^2+^ exchanger activity and undesirable EADs. For instance, ATX-II was shown to promote atrial fibrillation ^2^. Using this artifice, drugs such Ranolazine and GS458967 ^3^, as well as a set of natural compounds extracted from plants, have been identified that block I_NaL_ and that should represent good anti-arrhythmic drugs.

While the pathological contribution of I_NaL_ to arrythmia generation is now well documented, there are however several important issues that require clarification. Unquestionably, Na_v_1.5 is the predominant Na_v_ isoform expressed in the heart, but other subtypes have been detected as well, all capable to generate I_NaL_. In human cardiomyocytes derived from induced-pluripotent cells (hiPS-CM), the tetrodotoxin-sensitive (TTX-S) Na_v_1.1, Na_v_1.2, Na_v_1.3, Na_v_1.6 and Na_v_1.7 isoforms, as well as TTX-resistant (TTX-R) Na_v_1.8 isoform, have been detected as well ^4^. In rabbit ventricular myocytes, Na_v_1.4 is the predominant TTX-sensitive isoform ^5^ and increases excitation-contraction coupling by reversing the Na^+^ - Ca^2+^ exchanger mode. In mouse ventricular myocytes, four TTX-S Na_v_ isoforms are detected (Na_v_1.1, Na_v_1.3, Na_v_1.4 and Na_v_1.6), that represent up to 23% of the current ^6^. Interestingly, Na_v_1.5 is located at the cell surface with high density at intercalated disc, but relatively absent from transverse (t)-tubular system where TTX-S Na_v_ isoforms are the predominant isoforms ^6-10^. These findings imply a role of TTX-S channels in AP t-tubular transmission and excitation-contraction coupling that is far more important than what their contribution to total Na^+^ influx would suggest ^11, 12^. In dog ventricular myocytes, the TTX-S current represents up to 44% of I_NaL_ in physiological conditions ^13^, further indicating that TTX-S channels may well be implicated in pathological prolonged QT interval. The finding that cardiac arrythmias are commonly observed in several neuronal and musculoskeletal diseases ^14^ further hint to a putative role of TTX-S channels that needs to be mechanistically dissected. Conversely, the proposal that TTX-S Na_v_ channels are not required for excitation-contraction coupling in rat ventricular myocytes in physiological conditions ^15^ are also an indication that they may represent safe therapeutic targets for arrythmia treatment. If one compiles the wealth of literature on cardiac TTX-S Na_v_ channels, the experimental evidence can be summarized as follows: i) the expression levels and subtypes of TTX-S Na_v_ channels vary depending on species and tissue localization, ii) a significant proportion of TTX-S channels are generally located in the t-tubules where excitation-contraction coupling takes place, iii) I_NaL_ can be supported by TTX-S Na_v_ channels, and iv) direct implication of TTX-S Na_v_ channels in arrhythmia generation has been demonstrated. To segregate among Na_v_1.5, the predominant TTX-R channel, and the TTX-S Na_v_ isoforms, we dispose mainly of two tools. First, TTX that up to 100 nM will selectively block TTX-S channels and leave intact Na_v_1.5 contribution. For the time being selective blockers of each TTX-S Na_v_ isoform are lacking. Second, ATX-II an FDA-advocated I_NaL_ inducer but that is used in ill-defined conditions since the selectively profile of this compound is poorly defined. Yet, this is an essential issue as it is of importance to know whether the I_NaL_ current triggered by ATX-II is solely mediated by Na_v_1.5 or also by the TTX-S Na_v_ isoforms. These considerations indicate that selective activators of Na_v_1.5, not influencing TTX-S channels, and conversely selective activators of TTX-S channels would be beneficial to better demonstrate the potential arrhythmogenic importance of each type of Na_v_ player.

In this report, we define with precision and in a standardized manner the selectivity profile of ATX-II. We also identified AaH-II, a natural peptide from *Androctonus australis Hector* scorpion venom, as an interesting high affinity and selective promoter of cardiac TTX-S I_NaL_. We illustrate that preferential activation of TTX-S I_NaL_ triggers arrythmias in conditions where Na_v_1.5-mediated I_NaL_ is negligible. We advocate the use of AaH-II as an important additional tool that complements ATX-II to investigate the pathological contributions of cardiac I_NaL_ and to facilitate the identification of compounds that block I_NaL_ selectively originating from TTX-S Na_v_ isoforms, most of them as likely as Na_v_1.5 channels to contribute to arrhythmias.

## Materials and Methods

### Peptides and compounds

All the peptides used in this study (AaH-II, AaH-II^*^ and ATX-II) were synthesized by Smartox Biotechnology (https://www.smartox-biotech.com/). AaH-II* is a single amino-acid substituted analog of AaH-II (R^62^K mutation), previously developed to produce a photoactivable Na_v_ agonist ^16^. As shown on representative hNa_v_1.2 currents, AaH-II and AaH-II* are fully similar in their properties for inducing I_NaL_ (**Figure S1**). Interestingly, however, AaH-II* differs from AaH-II in lacking the inhibitory effect of AaH-II at the highest concentrations, which is presumably due to a second binding site on voltage-sensor domain I that we presume lost upon this R to K substitution ^17^. TTX was purchased from Tocris Bioscience.

### Ethical statement

All animal care and experimental procedures were performed in the animal facilities of l’institut du thorax that have been accredited by the French Ministry of Agriculture. All animals were exposed to 12 hr light/dark cycles (light, 8:00 AM to 20:00 PM) in a thermostatically controlled room with free access to food and water. The experimental procedures were approved by the regional ethic committees (CEEA-006 Pays de la Loire, France) and authorized by the French Ministry of National Education (APAFIS #34541-2022010310194375), Higher Education and Research according to the Directive 2010/63/EU of the European Union.

### Cell Culture

HEK293 cells stably expressing human (h)Na_v_1.1 (NM_001165963.4), hNa_v_1.2 (NM_001040142.2), hNa_v_1.3 (NM_001081676.2), hNa_v_1.5 (NM_000335.5), hNa_v_1.6 (NM_001177984.3) and CHO cells stably expressing Na_v_1.7 (NM_001365536.1) were cultured in Dulbecco’s Modified Eagle’s Medium (DMEM) supplemented with 10% fetal bovine serum, 1 mM pyruvic acid, 4.5 g/L glucose, 4 mM glutamine, 10 U/mL penicillin and 10 μg/mL streptomycin (Gibco, Grand Island, NY). HEK293 cells stably expressing human Na_v_1.4 (NM_000334.4) were cultured in Minimum Essential Medium (MEM) supplemented with 10% fetal bovine serum, 1 mM pyruvic acid, 4.5 g/L glucose, 4 mM glutamine, 10 U/mL penicillin and 10 μg/mL streptomycin (Gibco, Grand Island, NY). All cell lines were incubated at 37°C in a 5% CO_2_ atmosphere. G418 (400-800 µg/mL) was used to maintain stable expression of hNa_v_1.1, hNa_v_1.2, hNa_v_1.3, hNa_v_1.4, hNa_v_1.5 and hNa_v_1.6, whereas hygromycin B (200 µg/mL) was used to maintain hNa_v_1.7 expression. Cells prepared for electrophysiological experiments were firstly detached using trypsin and isolated cells were diluted (∼300,000 cells/mL) in medium containing (in mM): 140 NaCl, 4 KCl, 2 CaCl_2_, 1 MgCl_2_, 5 glucose, and 10 HEPES (pH 7.4, osmolarity 298 mOsm).

### High-throughput automated patch-clamp

Automated patch-clamp recordings were performed with the SyncroPatch 384PE from Nanion (München, Germany) to investigate efficacy and/or selectivity of AaH-II, AaH-II^*^, ATX-II and TTX on sodium currents. Chips with single-hole medium resistance of 4.51 ± 0.01 MΩ (n=384) were used for recordings. Pulse generation and data collection were performed with the PatchControl384 v1.5.2 software (Nanion) and the Biomek v1.0 interface (Beckman Coulter). Whole-cell recordings were conducted according to the recommended procedures of Nanion. Prior to recordings, dissociated cells were shaken at 200 RPM in a cell hotel reservoir at 10°C. After initiating the experiment, cell catch, seal and whole-cell formation, liquid application, recording, and data acquisition were all performed sequentially and automatically. For sodium channel recordings, intracellular solution was (in mM): 10 CsCl, 110 CsF, 10 NaCl, 10 EGTA, and 10 HEPES (pH 7.2, osmolarity 280 mOsm) and the extracellular solution contained (in mM): 100 NaCl, 4 KCl, 40 NMDG, 2 CaCl_2_, 1 MgCl_2_, 5 glucose and 10 HEPES (pH 7.4, osmolarity 298 mOsm). For sodium channel recordings, whole-cell experiments were performed at a holding potential of -100 mV at room temperature (20°C) and the sampling rate set at 20 kHz. Peptides were diluted in the extracellular solution containing 0.3% bovine serum albumin. Na^+^ currents were triggered at a test potential of 0 or +10 mV with a pulse every 5 s. The effects of each peptide on Na^+^ currents were measured at the end of a 10-min application time at room temperature (20°C).

### Patch-clamp on isolated neonatal cardiomyocytes

Single cardiomyocytes were isolated from the ventricles of neonatal C57BL/6J wild-type (WT) mice aged postnatal day 0 to 3. Neonatal hearts were rapidly excised and placed in Hank’s Balanced Salt Solution (HBSS; Gibco, Grand Island, NY) on ice. The atria were removed, and the ventricles were minced into small fragments to initiate mechanical dissociation. For enzymatic digestion, ventricular tissue fragments were transferred into 5 mL of digestion solution containing 0.714 mg/mL type II collagenase (Gibco, Grand Island, NY) and 0.1 mg/mL pancreatin in HBSS. Digestion was performed under gentle agitation at 37°C for 10 min. The supernatant, containing dissociated cells, was collected, and an equal volume of Dulbecco’s Modified Eagle’s Medium (DMEM) supplemented with 10% horse serum, 5% fetal bovine serum (FBS), 1% L-glutamine, 100 U/mL penicillin, and 100 μg/mL streptomycin was added to terminate enzymatic activity. This digestion process was repeated multiple times to maximize cell yield. Following enzymatic dissociation, pooled cell suspensions were centrifuged at 800 rpm for 10 min to pellet the cells. The resulting pellet was resuspended in the supplemented DMEM solution. To remove undissociated tissue fragments, the cell suspension was filtered through a 4-μm cell strainer. To minimize fibroblast contamination, cells were plated onto 60 mm-diameter Petri dishes and incubated at 37°C for 1.5 hrs, allowing differential adhesion of fibroblasts. Non-adherent cardiomyocytes were then collected, resuspended, and diluted to a final concentration of approximately 300,000 cells/mL in a physiological extracellular solution containing (in mM): 140 NaCl, 4 KCl, 2 CaCl2, 1 MgCl2, 5 glucose, and 10 HEPES (pH 7.4, osmolarity 298 mOsm). The resulting cardiomyocyte suspension was immediately used for high-throughput patch-clamp recordings.

### Calcium imaging in isolated cardiomyocytes

Eleven-week-old male Wistar-Han rats were euthanized by intraperitoneal (i.p.) injection of a lethal dose of pentobarbital (140 mg/kg, Euthasol®). Subsequently, the hearts were quickly excised and rinsed in cold calcium-free Tyrode solution with the following composition (in mM): 118 NaCl, 4.17 KCl, 1.2 MgSO_4_, 1.2 KH_2_PO_4_, 21 NaHCO3, 11 glucose, 5 taurine, at pH 7.4. The aorta was promptly cannulated, and the hearts perfused in a retrograde fashion with warm (37-38°C) Tyrode containing 1.25 mM CaCl_2_ to flush the coronary arteries of blood and next with a Tyrode solution supplemented with 1 mg/mL collagenase type 2 (Worthington Biochemical Corp, Lakewood, NJ) and 0.2 U.mL^-1^ protease type XIV (Sigma, Poole, UK) during 15-20 min. Once enzymatic digestion is completed, ventricles were shred into small pieces with surgical scissors in 0.5 mM Ca^2+^ Tyrode solution, filtered through a 200-µm mesh and next resuspended in 1 mM Ca^2+^ Tyrode solution for 1 hr at RT. Isolated cardiomyocytes were incubated in the dark with Fura-2-AM calcium probe (2 µM in 20% v/v DMSO; Life Technologies) for 15 min. Cardiomyocytes were then loaded on the IonOptix system (IonOptix Corporation, U.S.A.) and perfused with the warm (37°C) Tyrode solution. Cardiomyocytes were stimulated at 0.5 Hz using a MyoPacer field stimulator (20 V, 7-ms) to achieve steady state. Fura-2 fluorescence (F) at 510 nm was used to measure cytoplasmic Ca^2+^ concentration variation by exciting with UV light at 340 and 380 nm. Cells were paced for 1 to 2 min ensuring stabilization of the signals and the effect of the peptides were measured at the end of a 5- min application time. 30-sec-long periods were used as unit duration (representing 15 Ca^2+^ transients) during control and after the perfusion of the compound to count the number of arrhythmogenic transients.

### Ex vivo electrophysiological recordings

Eleven-week-old male Wistar-Han rats were first heparinized (heparin sodium, 0.5 U/g i.p.) and then euthanized by i.p. injection of 140 mg/kg pentobarbital (Euthasol®). Hearts were quickly excised and immersed in a cold modified Krebs-Henseleit solution (in mM): 116 NaCl, 5 KCl, 1.1 MgSO_4_, 7 H_2_O, 0.35 NaH_2_PO_4_, 27 NaHCO_3_, 10 glucose, pH 7.4. The aorta was cannulated and the hearts were perfused on a Langendorff system by warm (36.5-37.5°C) and oxygenated (95% 0_2_ / 5% CO_2_) Krebs-Henseleit solution supplemented with 1.8 mM CaCl_2_, 1 mM lactate and 0.2 mM pyruvate (control solution) at a constant flow rate of 12 mL/min. Once connected to the Langendorff perfusion system, hearts were perfused with control solution for 10 min to stabilize their activities. Electrical activity of the upper part of the right ventricle was assessed by using a 64-electrode array (0.2-mm electrode diameter; 0.36-mm interelectrode distance) connected to EMS64-USB-1002 amplifier. Single-lead ECG was acquired to monitor global cardiac electrical activity. Data were sampled at 10 kHz per channel and acquired by EMapRecord5 (Mappinglab, Oxford, UK). Data were obtained at baseline (2 min before compound perfusion) and during AaH-II^*^ perfusion for 10 min at various concentrations. Data were analyzed by using EMapScope5 (Mappinglab, Oxford, UK). Activation times were determined as the point of maximal negative slope. All activation times were related to the timing of the first detected waveform and then used to draw activation maps. An average of 3 complexes was used for activation time and conduction velocity analyses, while interbeat interval was measured as the average of 10 consecutive intervals. The index of inhomogeneity was calculated as previously described ^18^. Arrythmias were quantified over a period of 2 min in control and 10 minutes after peptide perfusion. Hearts exhibiting sustained (> 1 min) ventricular extrasystoles were classified as arrhythmic.

### Electrocardiography

Eleven-week-old Wistar-Han male rats were anesthetized with 1.5% isoflurane using a nose cone (0.3 L/min after 2-5 min induction of anesthesia in a chamber containing 4-5% isoflurane). Rectal temperature was monitored continuously and maintained at 37-38°C using a heat pad. Six-lead ECGs were recorded with 25-gauge subcutaneous electrodes on a computer through an analog-digital converter (IOX2 2.10.0.40, EMKA Technologies, France). Sampling rate is set to 1 kHz and ECG were recorded during 2 min as control and during 10 min after AaH-II* intravenous (i.v.) injection at 3, 15 or 30 µg/kg using a 100 µL injection volume. Quantification of ECG parameters was performed in lead I (ECG Auto v3.5.5.28, EMKA Technologies) and data from at least 10 consecutive complexes were averaged. Standard criteria were used for interval measurements. The end of the T wave was defined as the point at which the slow component returned to the isoelectric line. Area of T wave is measured between S wave and the end of T wave. QT intervals were corrected for heart rate using the formula, QTc=QT/(RR/100)^1/2 19^. Blood samples (1 mL) were collected and centrifuged to extract plasma aliquots (50 µL minimal volume). Samples were stored at -20°C to later determine circulating peptide concentration by chromatography-tandem mass spectrometry (LC-MS/MS)

### Liquid chromatography-tandem mass spectrometry

Circulating AaH-II^*^ and TTX concentrations were analyzed in plasma samples by LC-MS/MS. All solvents were LC-MS grade and purchased from Biosolve (Valkenswaard, Netherlands). Standard solutions of AaH-II^*^ or TTX were prepared and serially diluted in water to obtain 10X standard solutions ranging 1-100 nmol/L. Standard solutions (10 µL) were then added to free plasma (90 µL) to get final concentrations ranging 0.1-10 nmol/L. Standard and test samples were then deproteinized with 250 μL of acetonitrile. After centrifugation (10 min, 10,000 × g, 4°C), supernatants were dried under a gentle stream of nitrogen for 30 min at room temperature. Dried samples were reconstituted in 100 µL of ultrapure water and transferred to vials for LC-MS/MS analyses. LC-MS/MS analyses were performed on a Xevo® TQD Absolute mass spectrometer with an electrospray interface and an Acquity PREMIER® UPLC™ device (Waters Corporation, Milford, MA, USA). Samples (10 μL) were injected onto a CSH-PREMIER column (1.7 μm; 2.1 × 100 mm, Waters Corporation) held at 60°C, and AaH-II^*^ or TTX were separated with a linear gradient of mobile phase B (acetonitrile, 0.1% formic acid) in mobile phase A (water, 0.1% formic acid) at a flow rate of 350 μL/min. Mobile phase B was kept constant for 0.5 min at 1%, linearly increased from 1 to 95% for 3 min, kept constant for 0.5 min, returned to the initial condition over 1 min, and kept constant for 1 min before the next injection. Targeted compounds were then detected by the mass spectrometer with the electrospray interface operating in the positive ion mode (capillary voltage, 3 kV; desolvatation gas (N_2_) flow and temperature, 800 L/h and 450°C; source temperature, 120°C). The multiple reaction monitoring mode was applied for MS/MS detection: *m/z* 1203.6 → 1328.3 (cone: 30 V, collision: 30 eV) and *m/z* 1444.2 → 1804.7 (cone: 30 V, collision: 20 eV). Chromatographic peak areas constituted the detector responses. Standard solutions were used to plot calibration curves for quantification (linear regression, 1/x weighting, origin excluded).

### Statistics

Values are represented as mean ± SEM (standard error of the mean). We performed all statistical analyses using GraphPad Prism 8.4.2. Comparisons between control group and compound groups are performed using Wilcoxon rank-sum test. For repeated measures one-way ANOVA was performed to compare multiple groups among various conditions followed by a Pukey post-test. A p-value lower than 0.05 was considered statistically significant.

## Results

### AaH-II^*^ is a peptide that preferentially activates TTX-sensitive cardiac Na_v_ channels

AaH-II^*^ has been discovered before the cloning of the various Na_v_ isoforms was completed. Hence, it was mainly used as a radioactive tracer to characterize binding sites in various tissues and examine functional effects at neuromuscular junctions, but surprisingly a complete pharmacological characterization on various human (h) Na_v_ isoforms has never been performed according to a standardized and statistical meaningful protocol. We took advantage of an automated patch-clamp setup to perform this characterization on a large set of hNa_v_ isoforms (**Figure 1**). Complete dose-response curves were constructed for hNa_v_1.5 and compared to the dose-response curves for all TTX-sensitive Na_v_ isoforms (hNa_v_1.1, hNa_v_1.2, hNa_v_1.3, hNa_v_1.4, hNa_v_1.6 and hNa_v_1.7). As shown, 0.1 nM AaH-II^*^ is sufficient to trigger important I_NaL_ by hNa_v_1.1, hNa_v_1.2, hNa_v_1.3 and hNa_v_1.6, a concentration at which hNa_v_1.5 is spared (**Figure 1*a***). These TTX-sensitive hNa_v_ isoforms have thus greater sensitivity to low concentrations of this peptide than the major cardiac Na_v_ isoform. A higher concentration of 0.3 nM is required to detect a noticeable I_NaL_ mediated by hNa_v_1.5 and hNa_v_1.4, while hNa_v_1.7 remains spared. A detectable I_NaL_ was measurable for hNa_v_1.7 only at 1 nM. Complete dose-response curves were built for all hNa_v_ isoforms and compared to the dose-response curve of hNa_v_1.5 (**Figure 1*b***). hNa_v_1.1, hNa_v_1.2, hNa_v_1.3 and hNa_v_1.6 were all confirmed as being more sensitive to low AaH-II^*^ concentrations than the hNa_v_1.5 Isoform, suggesting that I_NaL_ from these TTX-S isoforms can be activated in isolation if these isoforms are present in cardiac tissue. In contrast, hNa_v_1.4 had equal sensitivity to AaH-II^*^ than hNa_v_1.5, while hNa_v_1.7 was significantly less sensitive, indicating that moderate concentrations of AaH-II^*^ (<1 to 3 nM), susceptible to mildly activate hNa_v_1.5, should spare hNa_v_1.7. The EC_50_ values that quantify the AaH-II^*^ dose required to half-maximally stimulate I_NaL_ are given in **Table 1**. Clearly, hNa_v_1.1 with an EC_50_ as low as 0.06 nM was by far the most sensitive isoform to AaH-II^*^. The following rank order of sensitivity to AaH-II^*^ was thus as follows: hNa_v_1.1 > hNa_v_1.2 > hNa_v_1.6 > hNa_v_1.3 >> hNa_v_1.5 = hNa_v_1.4 >> hNa_v_1.7. **Figure 1*c*** also quantifies the maximum reachable level of I_NaL_ by AaH-II^*^. As seen, maximally effective AaH-II^*^ concentrations provide the greatest maximal I_NaL_ stimulations for hNa_v_1.1 and hNa_v_1.5 (∼19-fold), while producing the lowest I_NaL_ stimulation for hNa_v_1.4 (∼6-fold).

**Table 1.**
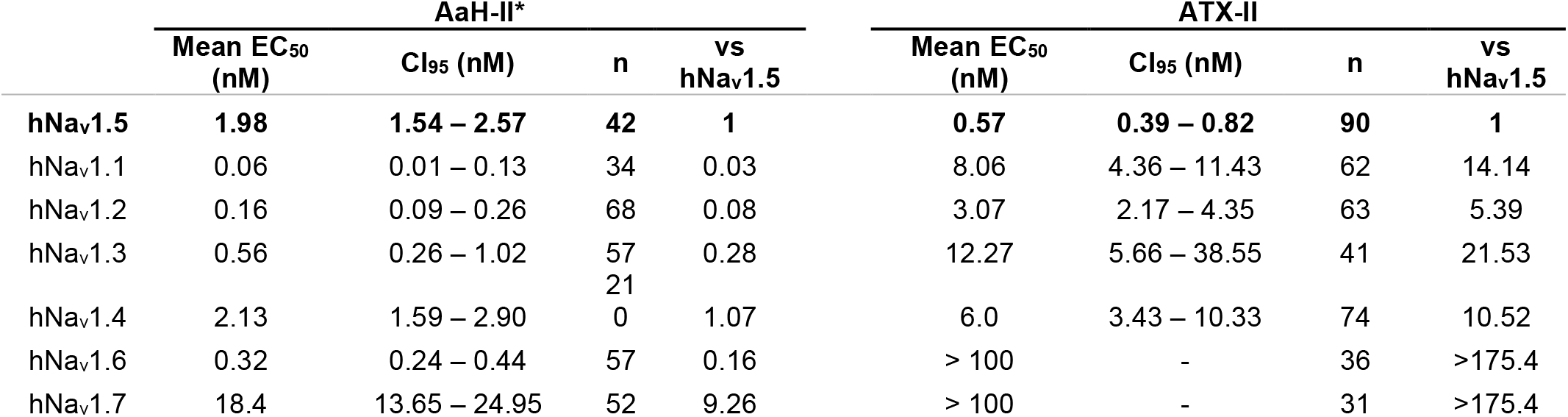
Comparison of AaH-II* and ATX-II selectivity among hNa_v_ subtypes. CI_95_: 95% confidence interval (n cells). Ratio of selectivity is calculated as hNa_V_1.x/hNa_V_1.5.

**Figure 1.**
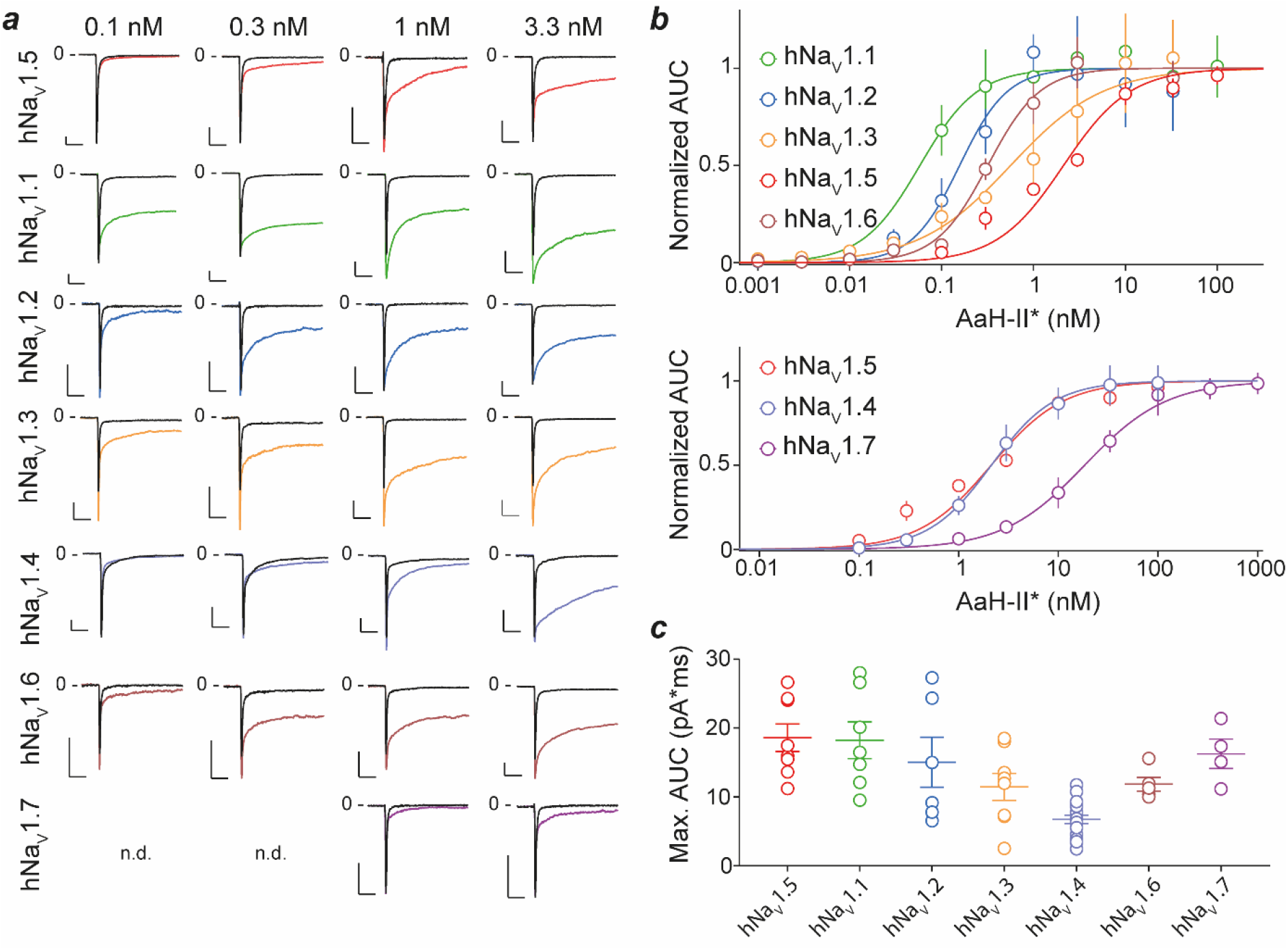
AaH-II* sensitivity of hNa_v_ channels. ***a***, Representative current traces elicited at a test potential of 0 mV for various hNa_v_ channels stably expressed in HEK293 cells (all except hNa_v_1.7) or CHO cells (hNa_v_1.7) before and after application of 0.1, 0.3, 1 or 3 nM AaH-II*. n.d.: not determined. Scale bars: all 0.5 nA and 10 ms. ***b***, Normalized average dose-response curves assessing the potency of AaH-II* on various hNa_v_ isoforms. IC_50_ values were calculated from the fit of the data and provided in **Table 1. *c***, Maximal I_NaL_ measured at highest effective AaH-II* concentration for each hNa_v_ isoform, as quantified by the Area Under the Curve (AUC). Mean ± SD values are shown.

For the purpose of comparison, we also examined the ATX-II sensitivity of these hNa_v_ isoforms examined in the same standardized conditions than those used for AaH-II^*^ (**Figure S2**). As shown, 0.1 to 0.3 nM ATX-II induce a significant I_NaL_ for hNa_v_1.5 that is not detected for TTX-sensitive hNa_v_ isoforms, except maybe marginally on hNa_v_1.2. According to the ATX-II dose-response curves performed, the EC_50_ value for inducing hNa_v_1.5 I_NaL_ was 0.57 nM (**Table 1**). The rank order of sensitivity to ATX-II was as follows: hNa_v_1.5 > hNa_v_1.2 > hNa_v_1.4 ≥ hNa_v_1.1 ≥ hNa_v_1.3 >>> hNa_v_1.6 = hNa_v_1.7. We found no evidence that ATX-II affects hNa_v_1.6 or hNa_v_1.7. In addition, the maximal reachable I_NaL_ was significantly lower for hNa_v_1.3 and hNa_v_1.4 compared to hNa_v_1.1 and hNa_v_1.2 (**Figure S2**). The rank order of sensitivity and the lack of sensitivity of hNa_v_1.6 and hNa_v_1.7 are therefore inverse to those observed for AaH-II^*^ indicating that ATX-II can be used for preferential activation of Na_v_1.5 I_NaL_ in cardiac tissues at 0.1 to 0.3 nM, while AaH-II^*^ is an indicated pharmacological tool for the preferential activation of several TTX-sensitive I_NaL_ at 0.1 nM.

### AaH-II^*^ pharmacological effect does not preclude TTX-sensitivity

Because TTX will be used to assess the importance of TTX-sensitive I_NaL_ in cardiac arrythmias, we made sure that AaH-II^*^-modified currents were still sensitive to 100 nM TTX. As shown, addition of 10 nM AaH-II^*^ to hNa_v_1.6-expressing cells induces an important uprise in I_NaL_ (**Figure S3*a-c***). A further addition of 100 nM TTX leads to a complete block of hNa_v_1.6 currents indicating, as expected ^17, 20^, that the binding sites for these two pharmacological agents do not overlap, nor that an allosteric competition takes place. The opposite experiment was also performed wherein hNa_v_1.6 currents were first fully blocked by 100 nM TTX and the subsequent application of 10 nM AaH-II* fails to activate any form of current, including I_NaL_ (**Figure S3*d-f***). We next investigated whether this finding also applies to freshy dissociated neonatal mouse cardiomyocytes. Total Na_v_ Na^+^ currents were recorded by automated high-throughput patch-clamp at -20 mV. As shown, 100 nM TTX leads to about 14.1 ± 2.1 % current block, indicating the existence of a minor proportion of TTX-sensitive channels in these cells (**Figure 2*a***,***b***). Further addition of 0.1 nM AaH-II*, that does not affect Na_v_1.5 channels, has no effect on current amplitude or I_NaL_ measured by the AUC. Conversely, if 0.1 nM AaH-II* is applied first, the current amplitude is significantly increased, as measured both by peak amplitude or AUC (**Figure 2*c***,***d***). Subsequent addition of 100 nM TTX largely reverses this current increase, indicating that 0.1 nM AaH-II* affects only TTX-sensitive Na_v_ channels in these mouse neonatal ventricular cardiomyocytes.

**Figure 2.**
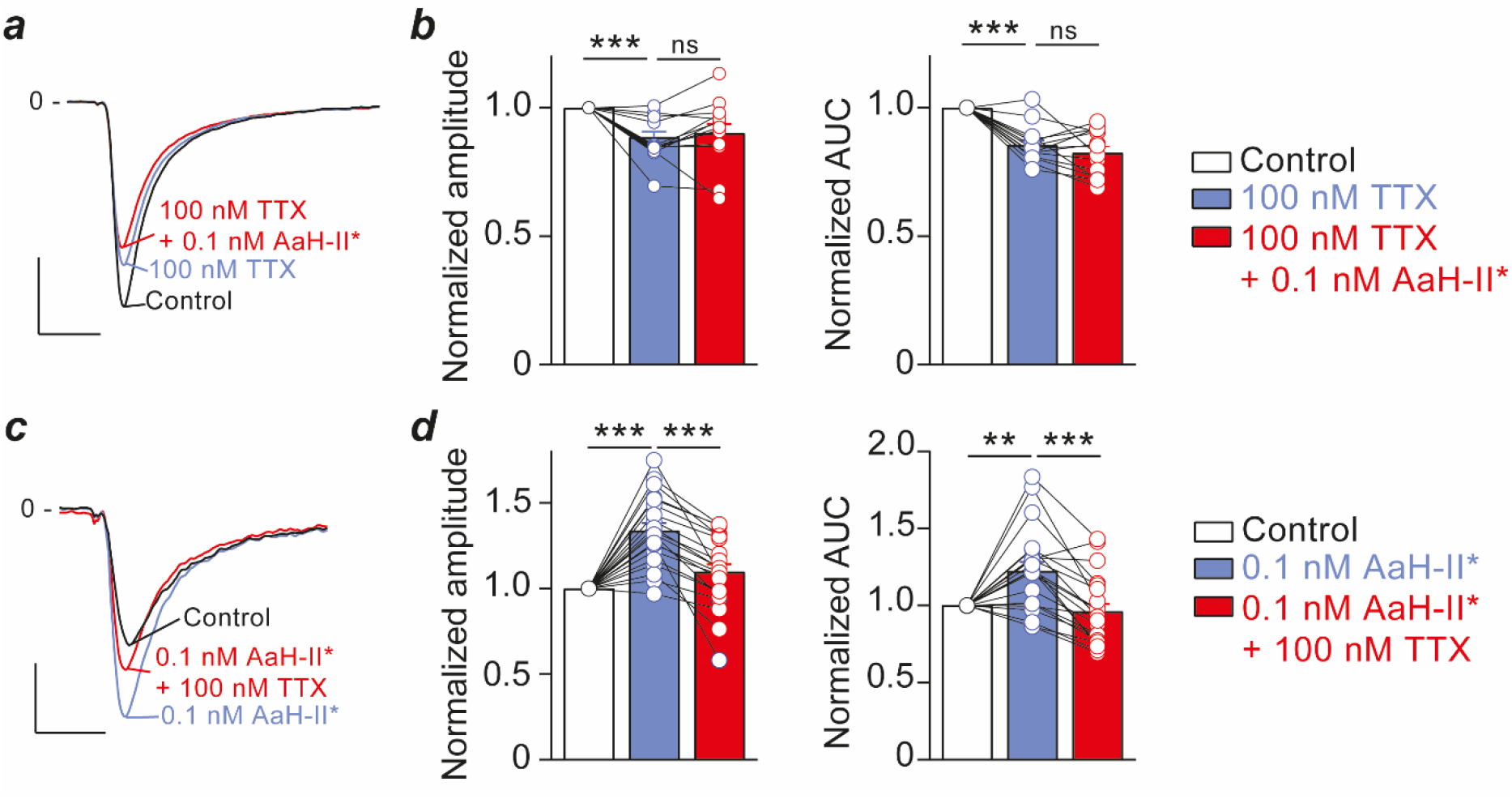
AaH-II* stimulates TTX-sensitive Na_v_ channels in neonatal mouse cardiomyocytes. ***a***, Representative Na^+^ currents from neonatal cardiomyocytes (holding potential -120 mV, test potential -20 mV) before and after application of 100 nM TTX (first) and 100 nM TTX + 0.1 nM AaH-II* (second). Scale bars: 2 nA, 2 ms. ***b***, Average current amplitude (left panel) or AUC (right panel) examining the pharmacological treatments normalized to control condition (n=14 cardiomyocytes). ***c***, Representative Na^+^ currents from neonatal cardiomyocytes before and after application of 0.1 nM AaH-II* (first) and 100 nM TTX + 0.1 nM AaH-II* (second). Scale bars: 0.5 nA, 2 ms. ***d***, same as ***b*** but pharmacological treatment of AaH-II* first, followed by the combined TTX / AaH-II* treatment (n=21 cardiomyocytes). **, p<0.01; ***, p<0.001.

*Dose-dependent arrythmia induction by AaH-II* and TTX-sensitivity of the response* – Freshly dissociated ventricular cardiomyocytes from adult rats were prepared and electrically stimulated at 0.5 Hz to investigate the shape of spontaneous calcium release (SCR) events with the IonOptix system in various experimental conditions. We first confirmed the proarrhythmic effects of ATX-II. At 0.1 nM, it should induce I_NaL_ selectively from the Na_v_1.5 isoform. In these conditions, the peptide triggers the appearance of arhythmic SCR events in 3 out of 11 cells and with a frequency above 15 events per min (**Figure S4*a-c***). In addition, the amplitudes of the calcium transients were reduced, while their time to peak increased (**Figure S4*d***). Both the percentage of cells displaying arhythmic SCR events and the frequency of these events increase with higher ATX-II concentrations (1 and 3 nM). These observations are coherent with the fact that higher ATX-II concentrations produce greater I_NaL_ from Na_v_1.5 with 3 nM ATX-II producing sub-maximal effect (**Figure S2**). However, 3 nM ATX-II is also a concentration at which several TTX-sensitive Na_v_ isoforms display I_NaL_ (hNa_v_1.1, hNa_v_1.2, hNa_v_1.3 and hNa_v_1.4) suggesting that their participation to enhanced arhythmic behavior of the cardiomyocytes cannot be ruled out. To test the potential involvement of TTX-sensitive Na_v_ isoforms in arhythmic behavior of cardiomyocytes, we also tested the appearance of SCR events upon application of various concentrations of AaH-II^*^. At 0.1 nM AaH-II*, a concentration that should selectively trigger I_NaL_ from TTX-sensitive Na_v_ channels and spare Na_v_1.5, SCR events are also observed in 13 out of 23 cells and with an average frequency of 14 SCR/min (**Figure 3*a-c***). Increasing AaH-II* concentration to 1 or 3 nM further enhances the probability of the arhythmic events, probably both by enhancing the TTX-sensitive I_naL_ amplitude and by recruiting the Na_v_1.5 I_NaL_ component (**Figure 3*a-c***). These higher concentrations of AaH-II* slightly reduce the amplitude of the calcium transients and slow the time to peak (**Figure 3*d***). This first set of data indicates that the selective activation of TTX-sensitive Na_v_ channels does trigger arrhythmia in the absence of Na_v_1.5 contribution. A further demonstration of the implication of TTX-sensitive Na_v_ isoforms to arrhythmias can be achieved by blocking TTX-sensitive Na_v_ channels by mild TTX concentrations (100 nM) that spare Na_v_1.5. As shown, 100 nM TTX does by itself not trigger the appearance of SCR events (**Figure 3*e***). It does however reduce the amplitude and increase the time to peak of regular calcium transients moderately. These results are coherent with a tight coupling between TTX-sensitive Na_v_ channels and the Ca^2+^ mobilization machinery. In the presence of 100 nM TTX, 0.1 nM AaH-II^*^ triggers SCR events with a much lower frequency than in the absence of TTX with 2 responding cells over 17 (**Figure 3*g-i***). These results confirm that I_NaL_ from TTX-sensitive Na_v_ isoforms has the potential to trigger arrhythmia. They are coherent with the almost complete lack of contribution of Na_v_1.5 to I_NaL_ at this AaH-II^*^ concentration (**Figure 1**). Conversely, at 1 nM AaH-II*, a concentration that leads to the concomitant activation of TTX-sensitive and Na_v_1.5 I_NaL_, SCR events become far more frequent (**Figure 3*g-i***) in proportions that resemble what is observed in the absence of 100 nM TTX (**Figure 3*a-c***). As observed for TTX alone, the condition TTX + AaH-II* (0.1 or 1 nM) leads to decreased calcium transient amplitude and time to peak (**Figure 3*j***). Overall, these data demonstrate that I_NaL_ of cardiac TTX-sensitive Na_v_ isoforms are pro-arhythmic *in vitro* on isolated rat cardiomyocytes.

**Figure 3.**
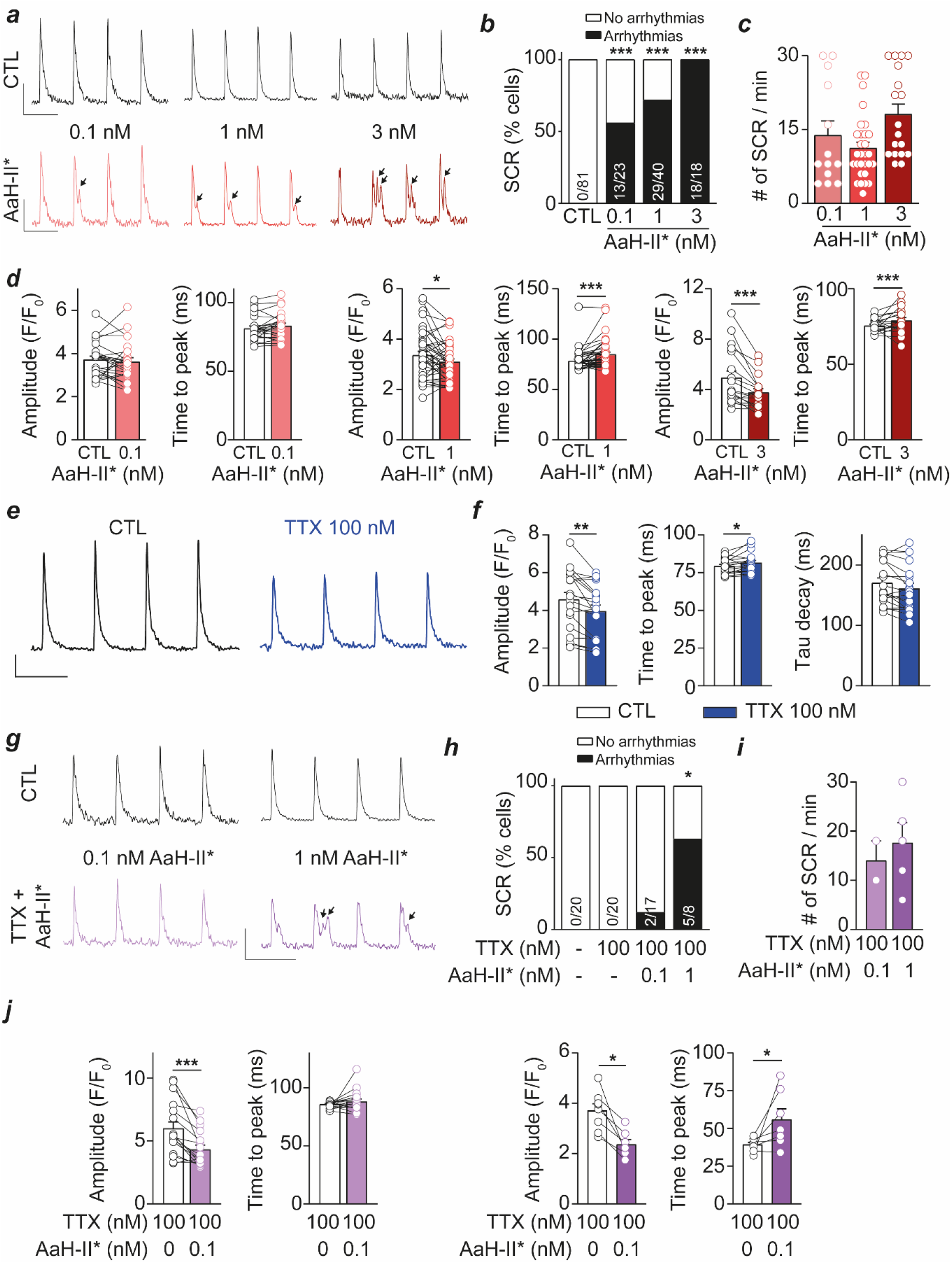
Effect of AaH-II* on calcium release from adult rat ventricular cardiomyocytes. ***a***, Calcium transients in control condition and after addition of 0.1, 1 or 3 nM AaH-II*. SCR events are shown by arrows. Scale bars: 1 F/F_0_; 2 sec. ***b***, Percentage of cells presenting SCR events in various AaH-II* concentration conditions. ***c***, Number of SCR events per min of recording as a function of AaH-II* concentration. ***d***, Quantification of the properties of elicited calcium transients in control condition and after application of 0.1, 1 or 3 nM AaH-II*. ***e***, Calcium transients in control condition and after perfusion of 100 nM TTX. Scale bars: 1 F/F_0_, 2 sec. ***f***, Quantification of amplitude, time to peak and tau decay of calcium transients in control condition and after perfusion of 100 nM TTX. ***g***, Representative calcium transients recorded in control condition and after application of 100 nM TTX. Scale bars: 1 F/F_0_; 2 sec. ***h***, Quantification of calcium transient properties before and after 100 nm TTX application. ***i***, Calcium transients in control condition and after combined treatment with 100 nM TTX and 0.1 or 1 nM AaH-II*. ***j***, Percentage of cells displaying SCR in various pharmacological conditions. ***i***, Number of SCR per min for cells that displayed SCR. ***j***, Properties of the calcium transients in the double pharmacological treatment condition (100 nM TTX + 0.1 or 1 nM AaH-II*).

### Ex vivo proarrhythmic effects of AaH-II* on TTX-sensitive Na_v_ channels

To check the proarrhythmic consequences of AaH-II* on cardiac tissue, we first evaluated the effects in isolated rat heart models to prevent confounding effects due to modulation of neuronal afferences in which TTX-sensitive channels are expressed. On the heart rhythm, measured by multielectrode array (MEA) on the upper part of the right ventricle (located near or on RVOT), 0.1 and 1 nM AaH-II* have no effect on heart rhythm, whereas 10 nM AaH-II* induces a significant ventricular bradycardia and heterogeneous repolarization (**Figure 4*a***,***b***). A rhythm reduction of this magnitude is expected to induce a global conduction block at the cardiac level. A significant proportion of this bradycardia is prevented by pretreatment with 100 nM TTX indicating the involvement of TTX-sensitive Na_v_ channels (**Figure 4*a***,***b***). The remaining bradycardia, that is still significant, appears to be due to the small bradycardic effect of TTX itself (**Figure S5*a***,***c***). 100 nM TTX is the concentration accepted that selectively blocks TTX-sensitive Na_v_ channels, without affecting Na_v_1.5^21^. This is indeed validated by the observation that perfusion of 100 nM TTX does not affect ventricular velocity supported by Na_v_1.5 (**Figure S5*b***,***d***)^22, 23^. Therefore, we conclude that the bradycardic effect of 10 nM AaH-II* requires the implication of TTX-sensitive Na_v_ channels as previously suggested^24^. At 10 nM AaH-II* we suppose that I_NaL_ is also carried by Na_v_1.5 in the cardiac tissue because this concentration is 100-fold higher than the concentration that spares Na_v_1.5 *in vitro*. These data are coherent with previous report indicating that I_NaL_ supported by gain of function mutations of Na_v_1.5 are not bradycardic ^19^. A closer examination of the shape of ventricular local field potential indicates that 0.1 nM AaH-II* significantly increases the repolarization area by 40.3 ± 8.0%. This effect is prevented by 100 nM TTX (**Figure 4*c***,***d***). This result validates the implication of TTX-sensitive I_NaL_ in the repolarization gradient of ventricular tissue. At 1 nM AaH-II*, the repolarization area is further increased to 140.6 ± 43.0%, an effect that is no longer prevented by 100 nM TTX (**Figure 4*c***,***d***). This later result indicates that perfusion of 1 nM AaH-II* leads to an increase of AaH-II* tissue concentration above 0.1 nM that is sufficient to increase I_NaL_ carried by Na_v_1.5. Therefore, the cardiac tissue should witness a concentration of AaH-II* in the vicinity of Na_v_1.5 that is between 0.1 and 1 nM. The increase of the area of repolarization by 0.1 nM AaH-II* leads to isolated premature ventricular contractions hearts studied and altered directionality of the electrical propagation but there are no sustained arrhythmias appearing (**Figure 4*e***,***f***). The further increase of the repolarization gradient at 1 nM AaH-II*, by the combined I_NaL_ activation from Na_v_1.5 and the augmented I_NaL_ carried by TTX-sensitive Na_v_ channels, leads to the emergence of sustained arrythmias in 5/7 hearts studied *ex vivo* (**Figure 4*e***,***f***). Pretreatment of the hearts with 100 nM TTX largely suppresses these arrhythmias and restores the directionality of the electrical propagation (**Figure 4*e***,***f***). The level of suppression is almost complete if 1 nM AaH-II* is used to promote arrhythmias. These results indicate that inhibition of TTX-sensitive Na_v_ channels helps preventing arrythmia triggered by the I_naL_ from TTX-sensitive channels and also from Na_v_1.5 if the extent of stimulation remains reasonable. However, if a larger activation of I_NaL_ carried by Na_v_1.5 is promoted by 10 nM AaH-II*, then the arrhythmogenic phenotype is only slightly decreased by 100 nM TTX pretreatment (from 6/7 arrhythmogenic hearts to 4/6; **Figure 4*e***). We next examined the impact of AaH-II* on cardiac conduction on the right ventricle. As shown, both 0.1 and 1 nM AaH-II* do not affect conduction (**Figure 4*g***,***h &* Figure S5**), whereas 10 nM AaH-II* significantly increases activation time (**Figure 4*g***,***h***). At 10 nM AaH-II* the effect is not prevented by 100 nM TTX indicating that Na_v_1.5 is the major contributor to conduction velocity, as shown previously^25^. It is also in agreement with the lack of effect of TTX on conduction velocity (**Figure S6**). In addition to slowing the conduction velocity, 10 nM AaH-II* increases the heterogeneity of the conduction (**Figure 4*j***). The absence of effect of 1 nM AaH-II* on conduction velocity is counterintuitive considering the effect of this concentration on repolarization gradient seen in **Figure 4*c***. We suppose that the level of I_NaL_ mediated by Na_v_1.5 is not high enough to affect conduction ^26, 27^.

**Figure 4.**
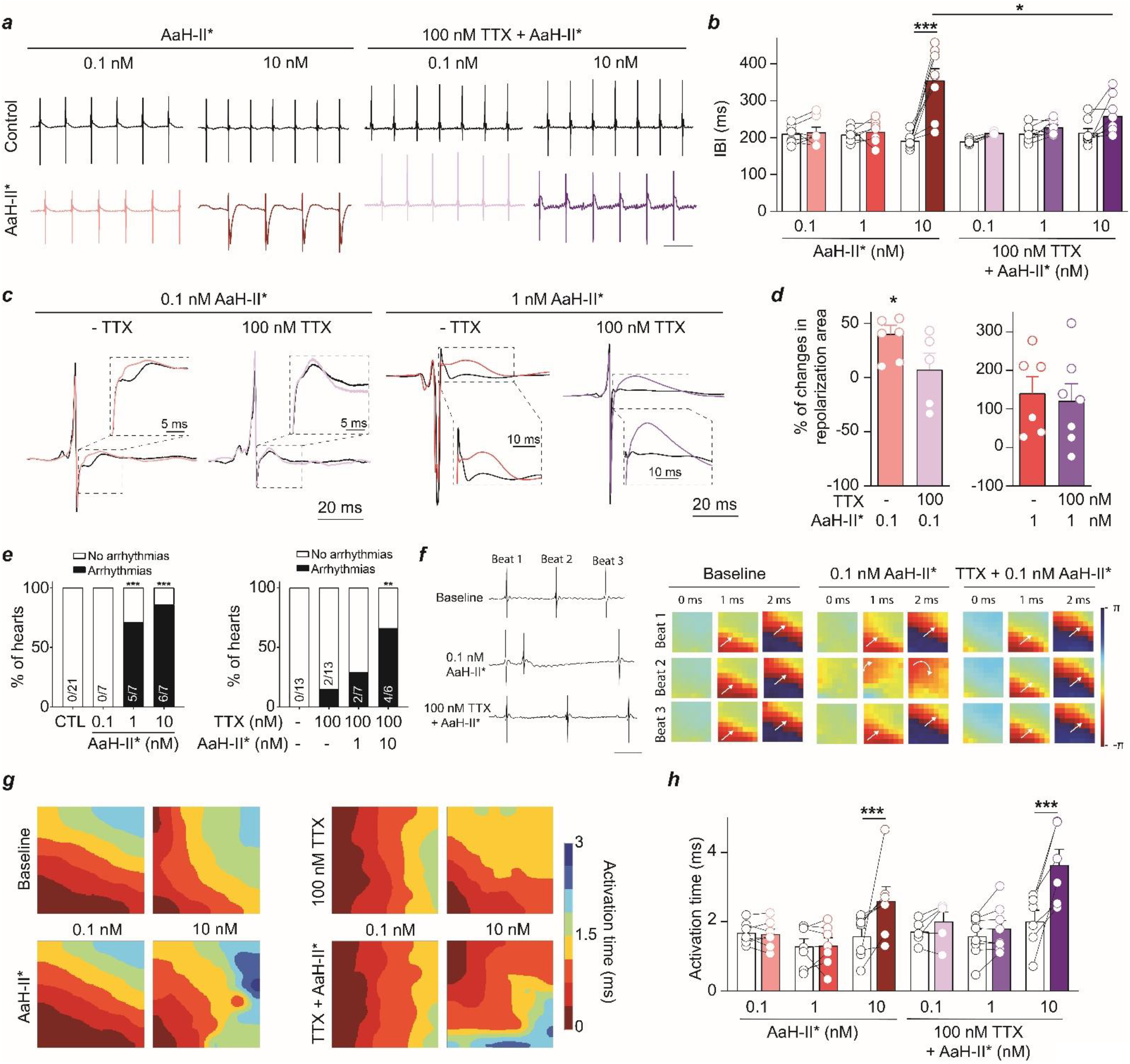
Arrhythmogenic effects of AaH-II* in isolated rat heart models. ***a***, Representative MEA recordings in control conditions and after perfusion of AaH-II* at various concentrations without and with pretreatment with 100 nM TTX. Scale bar: 250 ms. ***b***, Quantification of ventricular interbeat intervals (IBI) in various pharmacological conditions. *, *P*<0.05, ***, *P*<0.001. ***c***, Representative ventricular local field potentials at 0.1 and 1 nM AaH-II* without and with 100 nM TTX pretreatment. ***d***, Quantification of the percentage of increase in repolarization area in various pharmacological conditions. ***e***, Quantification of arrhythmogenic hearts in various pharmacological conditions. ***f***, Representative MEA recordings and phase maps highlighting the arrhythmogenic impact of AaH-II* and 100 nM TTX pretreatment on the directionality of the electrical propagation. The white arrow indicates this directionality. Scale bar: 100 ms. ***g***, Activation maps in various pharmacological conditions. ***h***, Quantification of activation time in various pharmacological conditions.

### In vivo proarrhythmogic effects of AaH-II*

Building on our *ex vivo* results, we investigated the *in vivo* effects of AaH-II* on rat heart activity. To this aim, we first investigated the plasmatic concentrations of AaH-II* upon intravenous injection (i.v.) of various doses of AaH-II*. 3 μg/kg AaH-II* leads to a circulating plasmatic concentration beneath 0.1 nM, i.e. the detection threshold of the mass spectrometry used (**Figure 5*a***). Increasing by 5- to 10-fold the dose of AaH-II* injected i.v. leads to plasmatic concentrations of 2.4 ± 0.4 nM (15 μg/kg) or 4.3 ± 0.6 nM (30 μg/kg), respectively. A plausible assumption is that the cardiac tissue concentration of AaH-II* is lower than its corresponding plasma concentration. In spite of these low concentrations, 3 μg/kg AaH-II* tend to produce a slight increase in RR interval and QTc value (**Figure 5*b***,***c***). These effects are largely augmented at 15 and 30 μg/kg AaH-II*. For instance, the RR interval increases from145.6 ± 2.5 ms to 181.6 ± 2.7 ms at 15 μg/kg which is believed to be due to AaH-II* effects on sympathetic innervation (**Figure 5*c***). This may also explain the increase in coefficient of variation for the RR interval. More interesting was the observation that QTc value increases from 46.1 ± 1.3 to 79.2 ± 4.6, which is supported by the increase in the T wave area from 1.2 ± 0.2 to 3.2 ± 0.5 mV/ms. This T wave area was further increased at doses higher than 15 μg/kg AaH-II* (**Figure 5*c***). Other ECG parameters are not affected (**Table 2**). At 3 μg/kg AaH-II*, none of the rats were arrhythmic, but all became arrhythmic if higher doses were used (**Figure 5*d***,***e***). At 15 μg/kg AaH-II*, atrio-ventricular (A-V) blocks and/or isolated PVCs are monitored in all rat. At 30 μg/kg AaH-II*, all rats presented arrhythmogenic events (A-V blocks/PVCs) with episodes of ventricular tachycardia (VT) / ventricular fibrillation (VF) in half of them. The latency of appearance of arrythmia was decreased upon increasing the dose of AaH-II* from 15 to 30 μg/kg (**Figure 5*f***). Next, we investigated how blocking TTX-sensitive Na_v_ channels would prevent these arrythmias. First, we evaluated the effect of various TTX i.v. doses from 3 to 243 μg/kg on ECG parameters (**Figure S7*a***,***b &* Table 2**). Consistent with the effect of TTX on *ex vivo* experiments (**Figure S6**), TTX increases the RR interval progressively with increasing doses. Interestingly, the QRS duration was not affected for doses up to 27 μg/kg, but suddenly increased at 81 μg/kg, indicating that doses above 27 μg/kg affect Na_v_1.5 activity. For these reasons, we decided to work at TTX dose equal to 27 μg/kg. This dose corresponds to a circulating TTX plasmatic concentration of 403 ± 44 nM (**Figure S7*c***). While this value is 4-fold higher than the concentration that should be used to be fully selective to TTX-sensitive Na_v_ channels, we also expect that intracardiac TTX concentration is lower in reality by the same order of magnitude. This concentration is also 25- to 75-fold lower than the one needed to block Na_v_1.5 (frequently 10 or 30 μM are used^21^). Importantly, rats pretreated with 27 μg/kg TTX are clearly non arrhythmic (**Figure 5*h***). In these experimental conditions, TTX prevented the appearance of VT / VF events at the various AaH-II* doses (**Figure 5*h***). Only A-V blocks and PVCs were monitored but with a clear reduction in the percentage of rats affected (**Figure 5*h***,***i***). Finally, in addition to preventing arrythmias, inhibition of TTX-sensitive Na_v_ channels normalized all ECG parameters with the exception of the RR interval (**Figure 5*j***). There is no longer a significant difference in RR interval between 3 and 30 μg/kg AaH-II* on the RR interval contrary to what was observed in the absence of TTX (**Figure 5*c***). We assume that TTX blocks the sympathetic effects of AaH-II*. The bradycardic effects of TTX alone are coherent with what is observed *ex vivo* (**Figure S6**) and is even more pronounced possibly due to the combination of an intrinsic cardiac and sympathetic effect. These results indicate that despite the fact that AaH-II* doses of 15 and 30 μg/kg potentially activate I_NaL_ from Na_v_1.5, inhibition of TTX-sensitive Na_v_ channels has remarkable anti-arrhythmic properties.

**Table 2.**
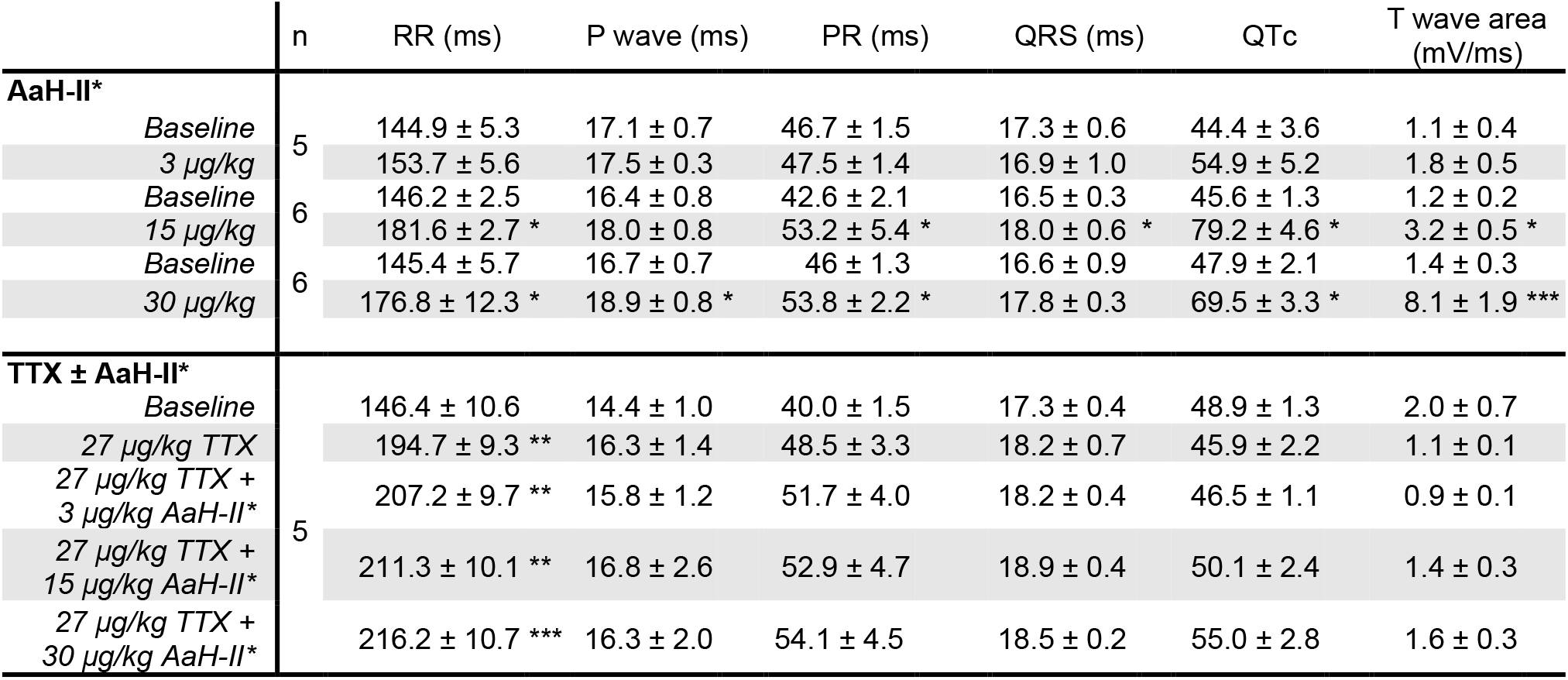
ECG parameters following AaH-II* or TTX + AaH-II* intravenous injection in anesthetized rats. n = number of rats. Wilcoxon matched-pairs signed rank test for AaH-II* only injection. Repeated Measures One-Way ANOVA followed by Holm-Sidak’s multiple comparisons test for TTX ± AaH-II* injection. p-value *p<0.05, **p<0.01, ***p<0.001 vs. Baseline. n = number of rats

|  | n | RR (ms) | P wave (ms) | PR (ms) | QRS (ms) | QTc | T wave area (mV/ms) |
| --- | --- | --- | --- | --- | --- | --- | --- |
| **AaH-II\*** |  |  |  |  |  |  |  |
| Baseline | 5 | 144.9 ± 5.3 | 17.1 ± 0.7 | 46.7 ± 1.5 | 17.3 ± 0.6 | 44.4 ± 3.6 | 1.1 ± 0.4 |
| 3 µg/kg |  | 153.7 ± 5.6 | 17.5 ± 0.3 | 47.5 ± 1.4 | 16.9 ± 1.0 | 54.9 ± 5.2 | 1.8 ± 0.5 |
| Baseline | 6 | 146.2 ± 2.5 | 16.4 ± 0.8 | 42.6 ± 2.1 | 16.5 ± 0.3 | 45.6 ± 1.3 | 1.2 ± 0.2 |
| 15 µg/kg |  | 181.6 ± 2.7 * | 18.0 ± 0.8 | 53.2 ± 5.4 * | 18.0 ± 0.6 * | 79.2 ± 4.6 * | 3.2 ± 0.5 * |
| Baseline | 6 | 145.4 ± 5.7 | 16.7 ± 0.7 | 46 ± 1.3 | 16.6 ± 0.9 | 47.9 ± 2.1 | 1.4 ± 0.3 |
| 30 µg/kg |  | 176.8 ± 12.3 * | 18.9 ± 0.8 * | 53.8 ± 2.2 * | 17.8 ± 0.3 | 69.5 ± 3.3 * | 8.1 ± 1.9 *** |
| **TTX ± AaH-II\*** |  |  |  |  |  |  |  |
| Baseline | 5 | 146.4 ± 10.6 | 14.4 ± 1.0 | 40.0 ± 1.5 | 17.3 ± 0.4 | 48.9 ± 1.3 | 2.0 ± 0.7 |
| 27 µg/kg TTX |  | 194.7 ± 9.3 ** | 16.3 ± 1.4 | 48.5 ± 3.3 | 18.2 ± 0.7 | 45.9 ± 2.2 | 1.1 ± 0.1 |
| 27 µg/kg TTX + 3 µg/kg AaH-II* |  | 207.2 ± 9.7 ** | 15.8 ± 1.2 | 51.7 ± 4.0 | 18.2 ± 0.4 | 46.5 ± 1.1 | 0.9 ± 0.1 |
| 27 µg/kg TTX + 15 µg/kg AaH-II* |  | 211.3 ± 10.1 ** | 16.8 ± 2.6 | 52.9 ± 4.7 | 18.9 ± 0.4 | 50.1 ± 2.4 | 1.4 ± 0.3 |
| 27 µg/kg TTX + 30 µg/kg AaH-II* |  | 216.2 ± 10.7 *** | 16.3 ± 2.0 | 54.1 ± 4.5 | 18.5 ± 0.2 | 55.0 ± 2.8 | 1.6 ± 0.3 |

**Figure 5.**
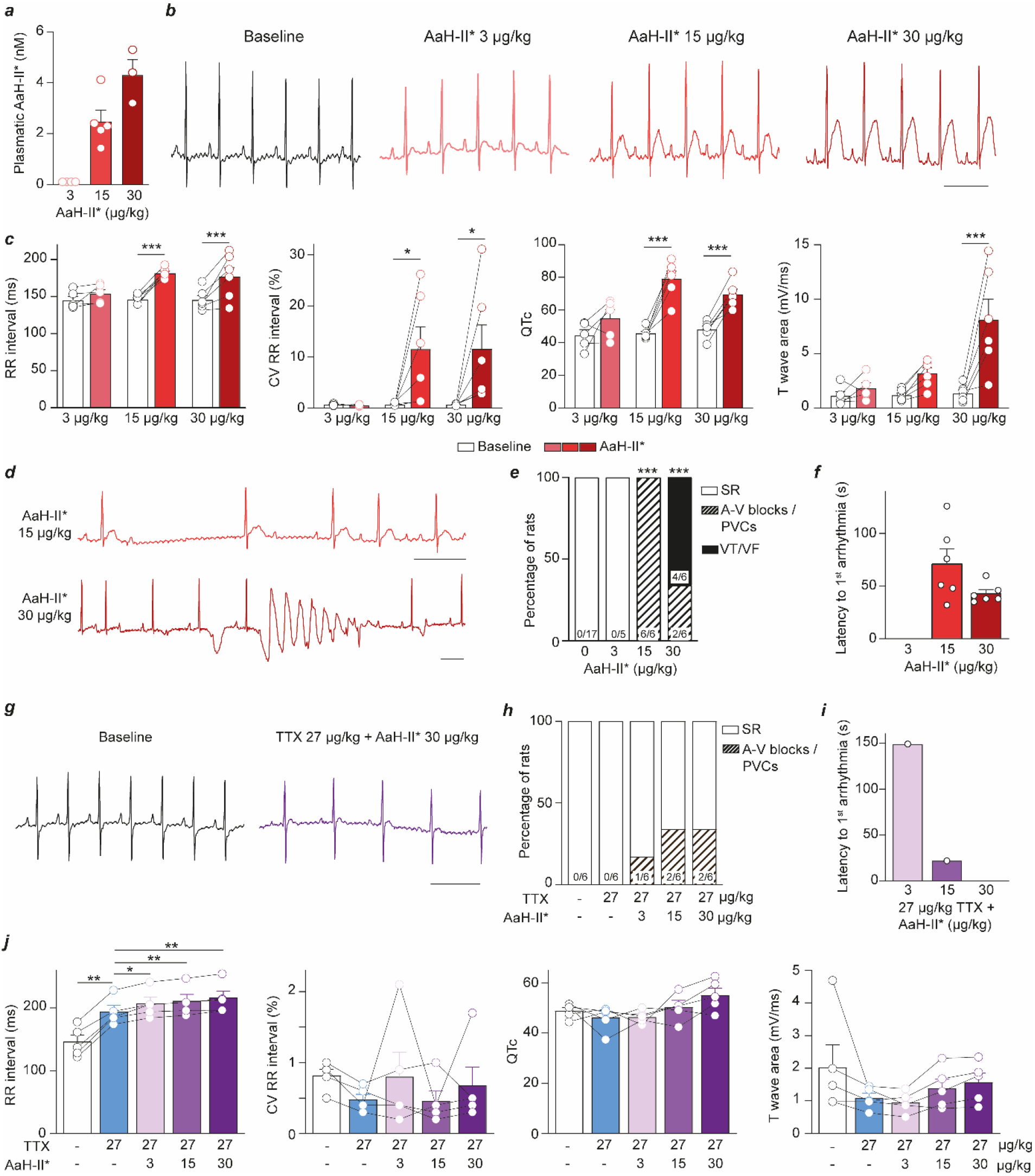
*In vivo* pro-arrhythmogenic effects of AaH-II*. ***a***, Circulating plasmatic concentrations of AaH-II* at three injected doses in rats. ***b***, Representative ECG recordings in control conditions or at three different doses of AaH-II*. Scale bar: 200 ms. ***c***, Quantification of ECG parameters at the three doses of AaH-II*. *, *P*<0.05. ***, *P*<0.001. ***d***, Representative recordings of A-V blocks and VT/VF at 15 and 30 μg/kg AaH-II*. ***e***, Percentage of arrhythmic rats and associated phenotypes. ***f***, Latency to first arrhythmia as a function of AaH-II* dose injected. ***g***, Effects of AaH-II on ECG parameters in rats pretreated with 27 μg/kg TTX. Scale bar = 200 ms. ***h***, Percentage of arrhythmic rats after TTX pretreatment. ***i***, Latency to first arrhythmia at various AaH-II* doses after TTX pretreatment. ***j***, Various ECH parameters for three AaH-II* doses on rats pretreated with 27 μg/kg TTX.

## Discussion

In this report, we examine the arrhythmogenic potential of I_NaL_ triggered by two different natural peptides, the well-known and widely used ATX-II and the less-well studied AaH-II*. We define the conditions of use of these two peptides with regard to their selectivity for the TTX-resistant hNa_v_1.5 channel versus the TTX-sensitive Na_v_ channels. We illustrate that a concentration of 0.1 nM allows ATX-II to be selective of hNa_v_1.5, while the same concentration of AaH-II* produces I_NaL_ only in TTX-sensitive Na_v_ channels. We also demonstrate that higher concentrations of ATX-II aggravate I_NaL_ from hNa_v_1.5 and recruits I_NaL_ from only two TTX-sensitive Na_v_ channels (hNa_v_1.1 and hNa_v_1.2). On the contrary, higher concentrations of AaH-II* has the advantage to moderately induce INaL from hNav1.5 while producing significant INaL values from the complete set of TTX-sensitive Na_v_ channels. These findings point to AaH-II* as being a much better cardiac I_NaL_ inducer than ATX-II, irrespective of the Na_v_ isoform being expressed in cardiac tissue and of the species under study. Careful use of AaH-II*, in defined concentration conditions, also allowed us to demonstrate that selectively triggering I_NaL_ from TTX-sensitive Na_v_ channels in the absence of any β-adrenergic stimulation, while sparring Na_v_1.5, is clearly proarrhythmic. The use of TTX confirmed the specificity of our selective I_NaL_ stimulation. Interestingly, arrythmias that occur in response to higher concentrations of AaH-II*, that produce a worsening of I_NaL_ from TTX-sensitive Na_v_ isoforms and a concomitant triggering of I_NaL_ from Na_v_1.5, are also largely reduced by the block of TTX-sensitive Na_v_ channels by 100 nM TTX. These findings have several important implications. First, they indicate that the arrhythmogenic potential of I_NaL_ from Na_v_1.5 are indirect and require the contribution of TTX-sensitive Na_v_ isoforms as relays for the coupling with the Ca^2+^ homeostasis machinery in the t-tubules. This would be coherent with the predominant localization of these TTX-sensitive Na_v_ channels in the t-tubules ^28, 29^. Second, they illustrate that examining INaL from cardiac TTX-sensitive Na_v_ channels is far more important in terms of cardiosafety than the I_NaL_ from Nav1.5. In that respect, we advocate the use of AaH-II* as a far more relevant drug to examine cardiac I_NaL_ in general than ATX-II. A shift in paradigm would be recommended. Third, it illustrates that a pharmacological inhibition of TTX-sensitive Na_v_ channels in the heart is a secure and efficient method to prevent arrhythmias triggered by I_NaL_, including if they arise from Na_v_1.5. Of course, considering the importance of TTX-sensitive Na_v_ channels in other organs, such an approach would be safe only if the action of the inhibitory drugs could be restricted to the heart.

How do these findings fit with earlier reports and extend their conclusions?

Our finding about the proarrhythmic potential of I_NaL_ supported by TTX-S Na_v_ channels is in line with earlier reports. In mice, Na_v_1.6 is present in t-tubules of cardiomyocytes and supports I_NaL_ triggered by the D96V mutation of calmodulin. It is of sufficient amplitude to promote aberrant Ca^2+^ release by RyR2 and prolonged AP duration which leads to LQT and ventricular tachycardia ^30^. In the same preparation, application of 4,9-ah-TTX, supposedly selective for blocking Na_v_1.6, appears to have antiarrhythmic effects on cellular arrhythmia triggered by the combined application of 4-aminopyridine and isoprenaline ^31^. The same compound actively reduces ATX-II promoted arrhythmias from calsequestrin-null ventricular cardiomyocytes. Also, in chronic heart failure and in atrial fibrillation (AF), Na_v_1.5 levels are reduced while TTX-sensitive Na_v_ channel expression are increases ^32, 33^. Under β-adrenergic stimulation and in mice models of AF (calsequestrin R33Q mutation), inhibition of TTX-sensitive Na_v_ channels by TTX or riluzole blocks aberrant Ca^2+^ oscillations, indicating that TTX-sensitive Na_v_ channels may represent interesting druggable targets for preventing AF ^34^. In yet another study, β-pompilidotoxin, a wasp venom peptide of 13 amino acids, was used to stimulate I_NaL_ from TTX-sensitive Na_v_ channels ^34^. The proarrhythmic nature of TTX-sensitive I_NaL_ could be revealed only in the presence of β-adrenergic stimulation. It is however unfortunate that this peptide requires a β-adrenergic agonist to reveal arrhythmias and possesses a rather poor affinity for its targets (reported EC_50_ of 21 μM on Na_v_1.2 compared to 0.16 nM for AaH-II* here; 131,200-fold better potency for AaH-II*) ^35^.

What does the future hold?

First, AaH-II* is a general I_NaL_-inducer from TTX-sensitive Na_v_ channels. While this is useful to replace ATX-II in assessing the proarrhythmic or protective potential of drugs acting on TTX-sensitive channels, it does not clearly indicate which isoform would be the best channel to target for developing antiarrhythmic drugs. Therefore, a large screening effort needs to be dedicated and pursued in identifying I_NaL_ inducers from each TTX-sensitive isoform. Part of this screening effort has been undertaken leading to the identification of JzTx-34 and Hm1a as I_NaL_ promoters from hNa_v_1.1 ^36, 37^.

Second, the finding that TTX-sensitive Na_v_ inhibitors are anti-arrhythmic, including in conditions where I_NaL_ is carried by Na_v_1.5, sparks interest in the anti-arrhythmic potential of these drugs in LQT3 animal models that carry a I_NaL_-promoting mutation on Na_v_1.5.

In conclusion, AaH-II* comes as a useful tool to discriminate the role of Na_v_1.5 and TTX-sensitive Na_v_ channels in arrhythmias. This tool will allow investigators to discriminate between drugs that act solely onto Na_v_1.5 or onto one of the Na_v_ components of TTX-sensitive channels.

## Notes

### Competing Interest Statement

The authors have declared no competing interest.

