## Supplemental data for "Late I_Na_ activation of cardiac TTX-sensitive sodium channels by AaH-II induces an arrhythmogenic phenotype"

*codirected this work.

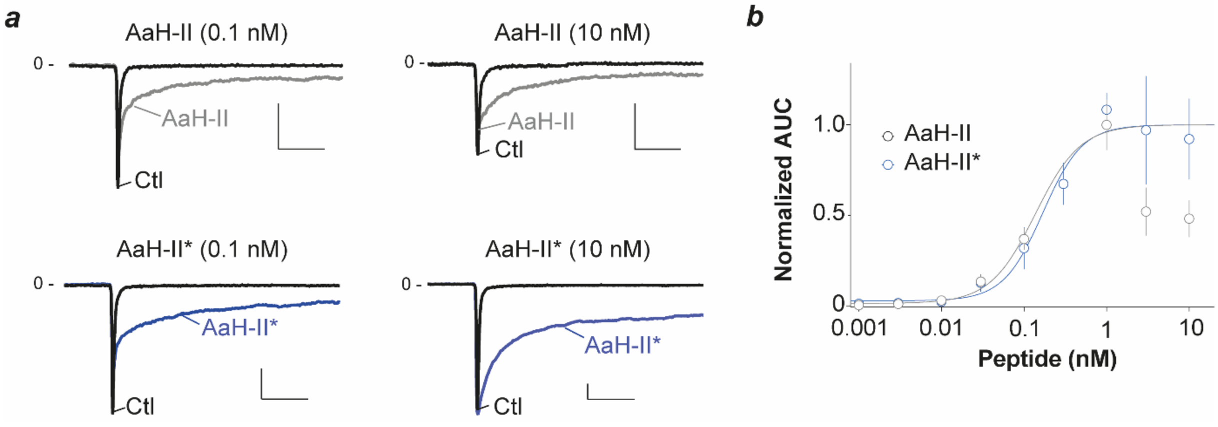


**Figure S1**. Pharmacological equivalence between AaH-II and AaH-II*. AaH-II* presents a R to K substitution at amino acid position 62. ***a***, Representative current traces of $hNa_v1.2$ before and after application of 0.1 or 10 nM AaH-II or AaH-II*. Scale bars: 0.5 nA and 10 ms. ***b***, Average normalized dose-response curves showing the effect of AaH-II and AaH-II* on the late $hNa_v1.2$ current (materialized by measuring AUC). Fit of the data yield $EC_{50}$ values of 0.11 nM (AaH-II, n=52 cells) and 0.16 nM (AaH-II*, n=57 cells), respectively. Please note that above 1 nM, AaH-II leads to peak current inhibition (see arrow in panel ***a***) that is not observed with AaH-II* and that is thought to occur by binding on a second lower affinity site on voltage sensor domain I [1].

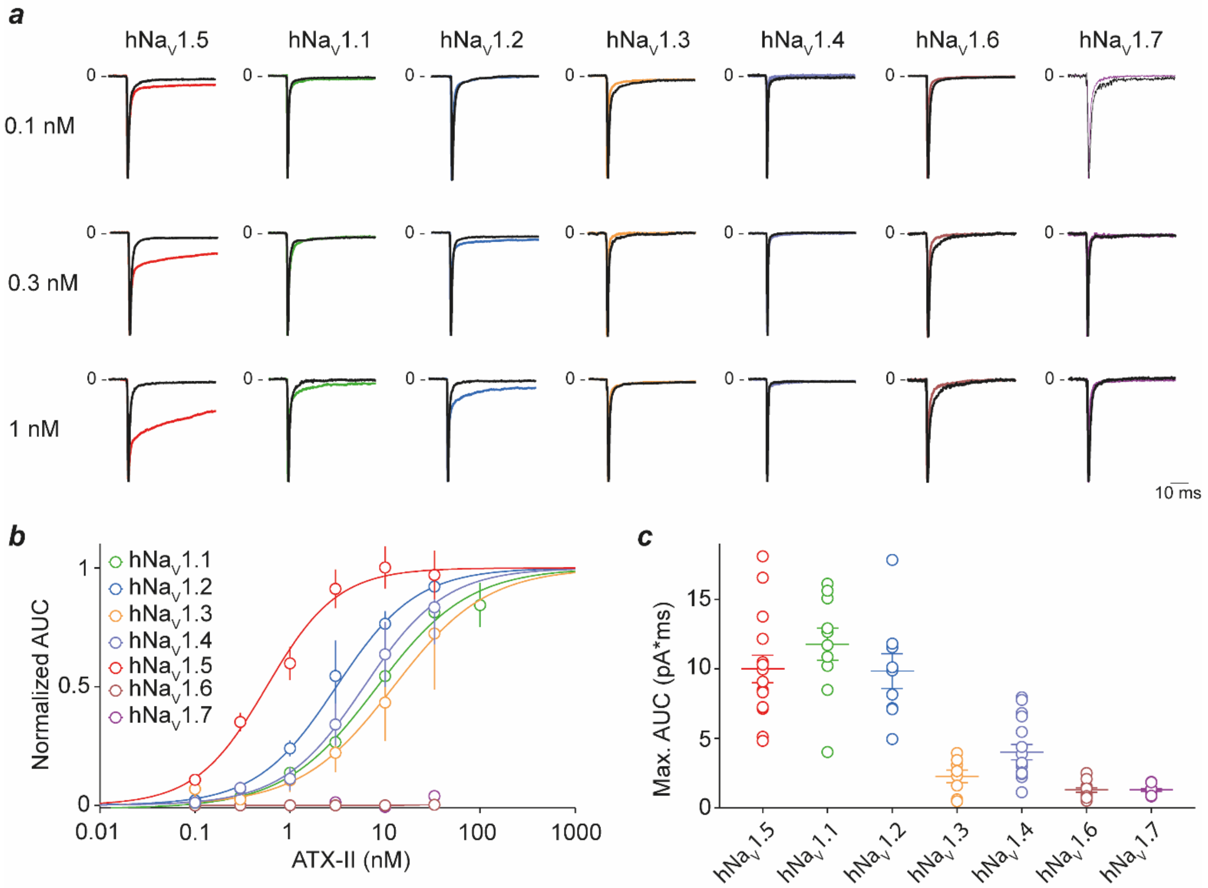


**Figure S2**. $I_{NaL}$ induction by ATX-II on various hNa$_v$ isoforms. ***a***, Representative normalized current traces elicited at test potential 0 mV before and after application of 0.1, 0.3 and 1 nM ATX-II. Note that 01 nM ATX-II only triggers $I_{NaL}$ from hNa$_v$1.5, and that 0.3 nM starts recruiting hNa$_v$1.2 $I_{NaL}$. ***b***, Average normalized dose-response curves illustrating the potency of ATX-II on various hNa$_v$ isoforms. $EC_{50}$ values obtained by the fit of the data are provided in **Table 1**. ***c***, Maximal level of $I_{NaL}$ triggered by ATX-II. hNa$_v$1.6 and hNa$_v$1.7 are insensitive to ATX-II action.

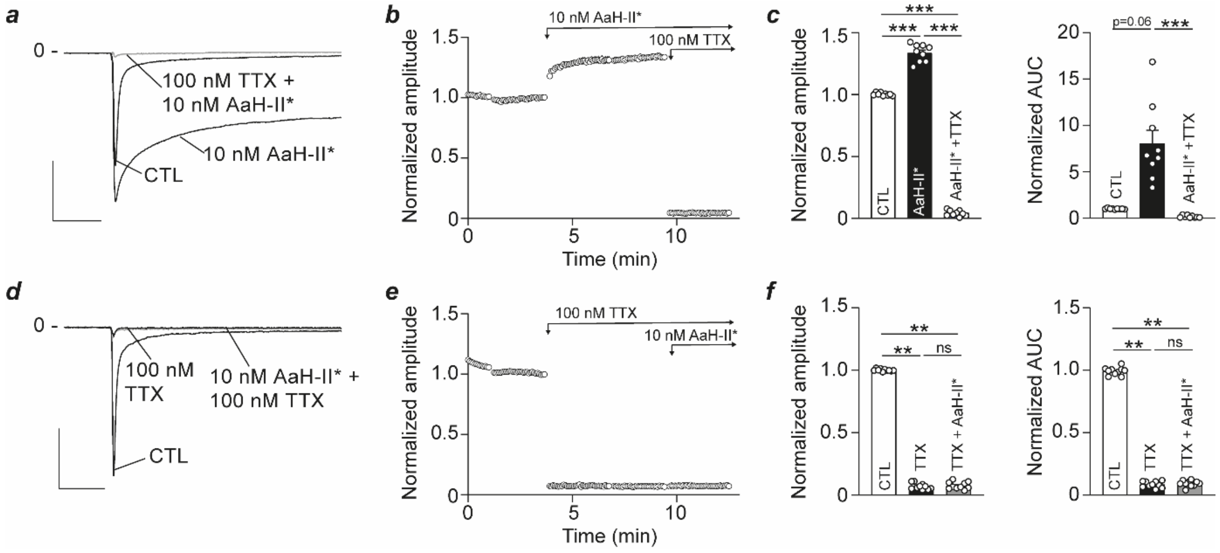


**Figure S3**. Preserved TTX sensitivity of AaH-II*-bound-hNa$_v$1.6. ***a***, Representative $Na^+$ current mediated by hNa$_v$1.6 at 0 mV test potential in control condition, after application of 10 nM AaH-II* to induce $I_{NaL}$, and after application of 100 nM TTX. Scale bars: 2 nA, 10 ms. ***b***, Representative time course illustrating the amplitude of hNa$_v$1.6 $Na^+$ currents before and during successive applications of 10 nM AaH-II* first, and 10 nM AaH-II* + 100 nM TTX after. ***c***, Average current amplitude (left) and AUC (right) illustrating the effect of AaH-II* and of TTX normalized to control condition (n=9 cells). ***d***, Representative $Na^+$ current mediated by hNa$_v$1.6 at 0 mV test potential in control condition, after application of 100 nM TTX to fully block the current, and after application of 100 nM TTX + 10 nM AaH-II*. Scale bars: 1 nA, 10 ms. ***e***, Representative time course illustrating the peak current amplitude of hNa$_v$1.6 $Na^+$ currents before and during successive applications of 100 nM TTX first, and 100 nM TTX + 10 nM AaH-II* after. ***f***, Average current amplitude illustrating the effect of 100 nM TTX and of 10 nM AaH-II* normalized to control condition (n=10 cells). ***, $p<0.001$.

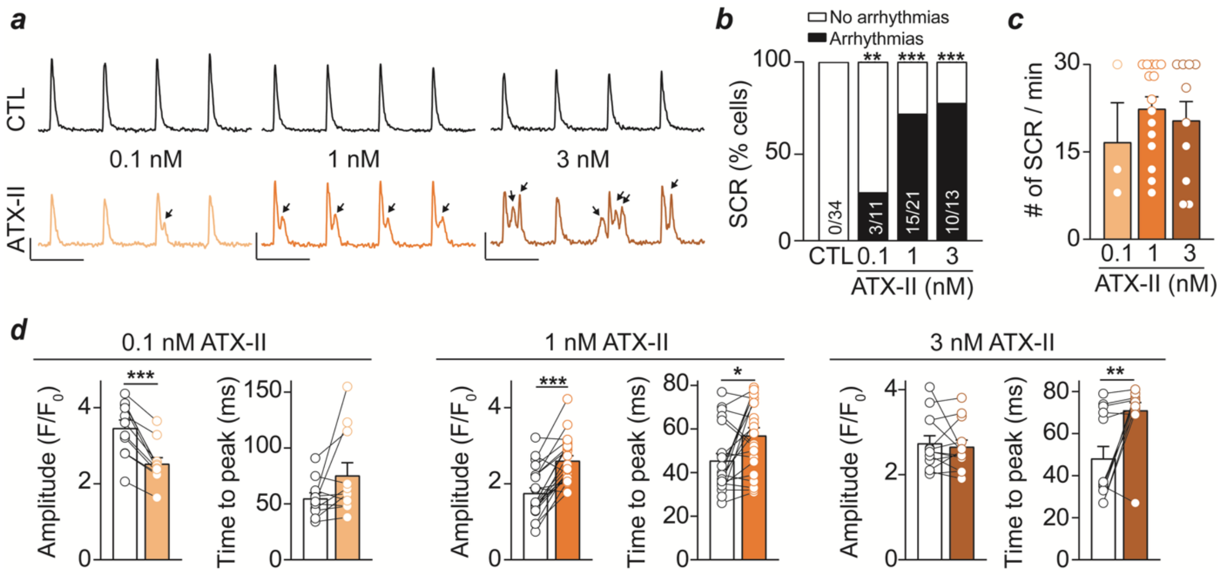


**Figure S4**. Proarrhythmic effects of ATX-II on adult rat ventricular cardiomyocytes. ***a***, Calcium transients triggered by electrical stimulation at 0.5 Hz in control conditions and after application of 0.1, 1 or 3 nM ATX-II. Abnormal spontaneous calcium release (SCR) events are indicated by arrows. Scale bars: 1 $F/F_0$, 2 sec. ***b***, Percentage of cells that display SCR at various ATX-II concentrations. ***c***, Number of SCR events per min of recording at various ATX-II concentrations. ***d***, Amplitude and time to peak quantification of normal calcium transients before and after 0.1, 1 or 3 nM ATX-II application.

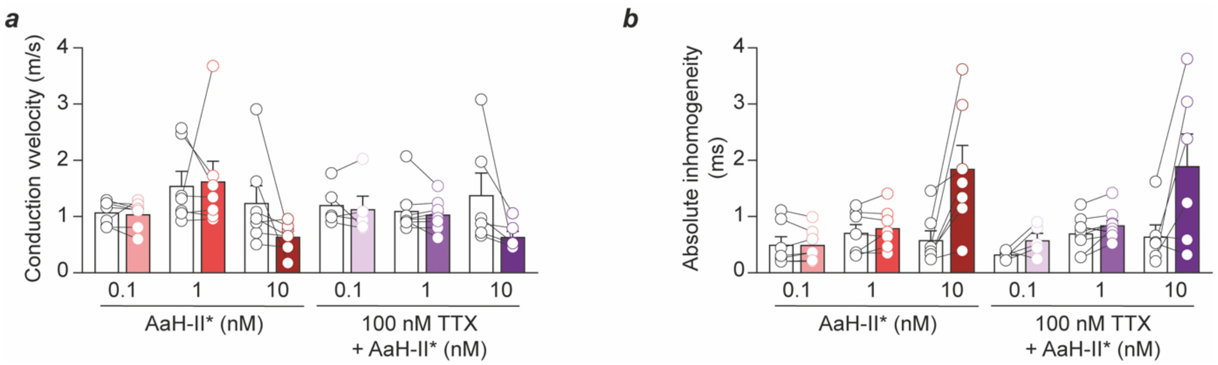


**Figure S5**. AaH-II* and TTX effects on conduction velocity parameters. ***a***, Quantification of conduction velocity in various pharmacological conditions. ***b***, Quantification of absolute inhomogeneity of conduction velocity in various pharmacological conditions.

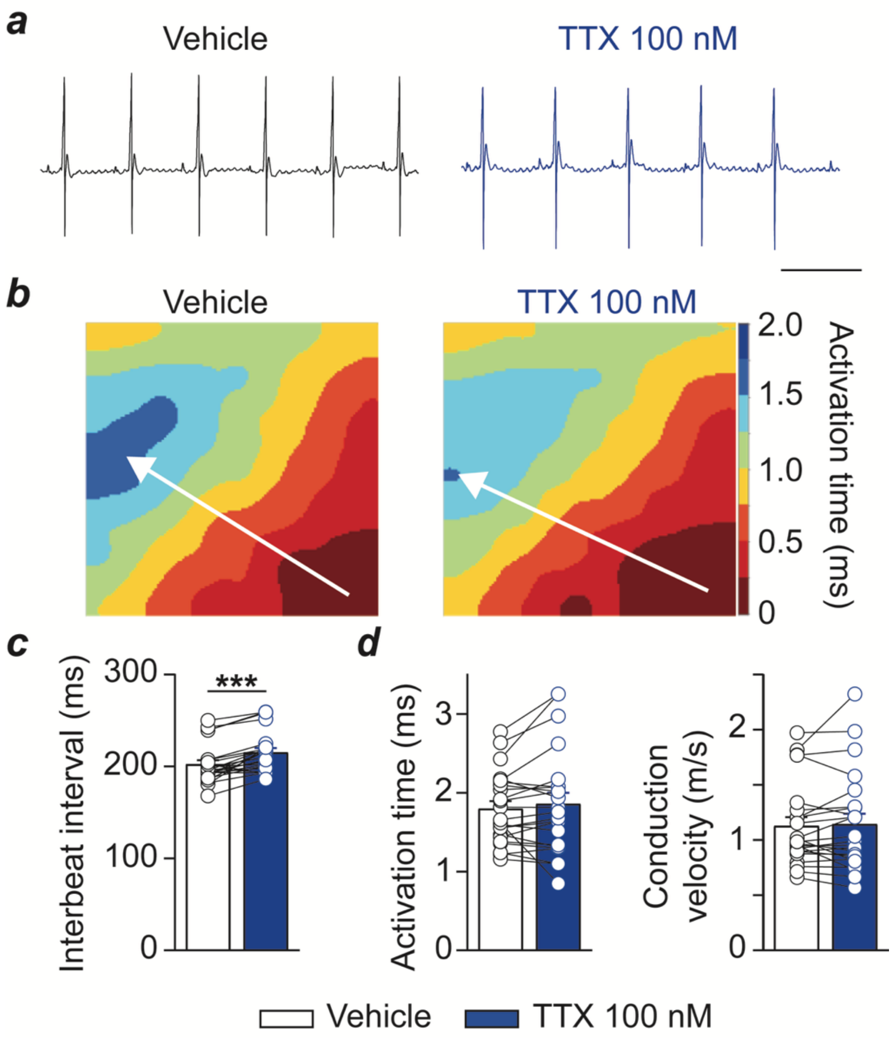


**Figure S6**. Bradycardic effect of TTX at a concentration that does not affect $Na_V1.5$. ***a***, Representative MEA recording in control and after perfusion of 100 nM TTX in isolated rat heart. Scale bar: 200 ms. ***b***, Resulting activation map. ***c***, Average interbeat interval and the modification induced by 100 nM TTX. ***, *P*<0.001. ***d***, Activation time and conduction velocity unaffected by 100 nM TTX.

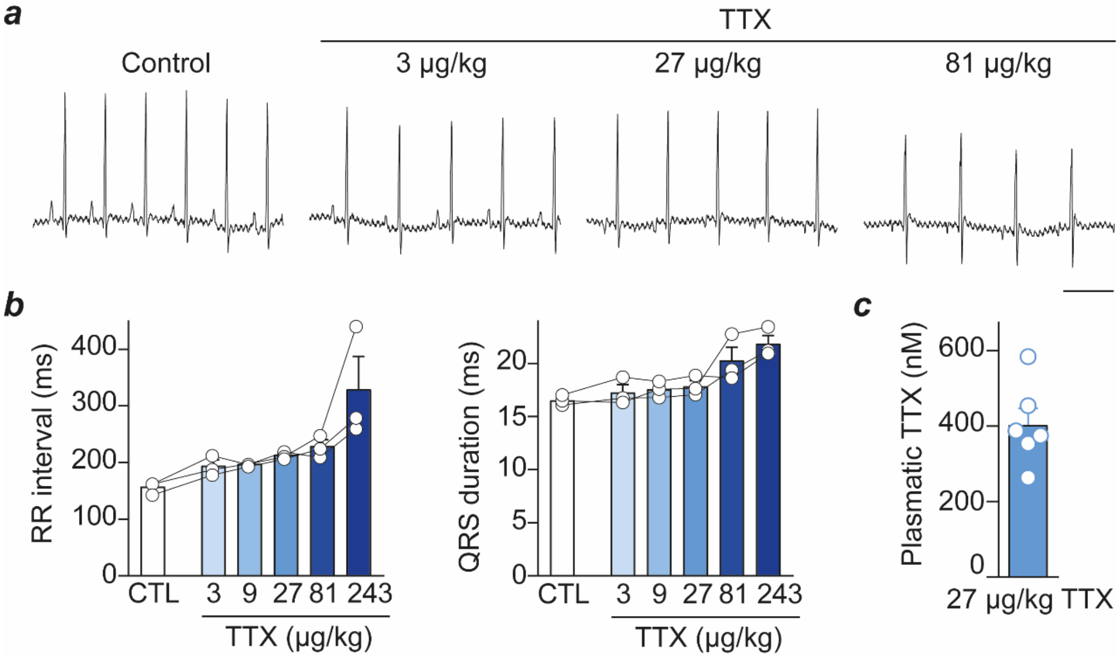


**Figure S7**. Effect of increasing *in vivo* doses of TTX on ECG parameters. Representative ECG recordings in control condition and at various doses of TTX. Scale bar: 200 ms. ***b***, Average RR interval and QRS duration at various conditions TTX doses compared to control condition. ***c***, Plasmatic concentration of TTX at the dose of 27 µg/kg.
